# Neural Bases for Working Memory of Natural Sound Textures

**DOI:** 10.64898/2026.10.28.762478

**Authors:** Yousef Mohammadi, Alexander J. Billig, Joel I. Berger, Maria Chait, Timothy D. Griffiths

## Abstract

Holding sound objects in mind is critical to understanding the sound world around us. Auditory texture perception is thought to rely on extracting statistical properties of sounds over time, raising the question of how encoding duration influences subsequent maintenance of auditory texture information. We investigated this question using behavioural performance and fMRI while participants performed two delayed match-to-sample tasks: a texture discrimination and a comparison loudness discrimination task. In the texture task, participants judged whether two sequentially presented sounds belonged to the same texture type, whereas in the loudness task, participants judged whether the sounds, drawn from the same texture type, had the same loudness. Stimuli were presented for either 0.1s or 2s. Discrimination sensitivity increased with duration in both tasks, but more strongly for texture discrimination. fMRI analysis revealed distinct effects of duration on univariate and multivariate activity in the auditory cortex and medial temporal lobe (MTL) during maintenance. In the auditory cortex, we found no univariate duration effect during maintenance, but multivariate patterns encoded texture category more strongly for longer stimuli, regardless of task. In contrast, MTL activity was modulated by sound duration in both tasks, but there was no effect of duration on texture category decoding. There was a task effect in MTL where multivariate patterns in posterior hippocampal and parahippocampal regions distinguished texture from loudness tasks following 2s stimuli. Together, these findings show effects of duration on sound texture representations during maintenance and that MTL representations reflect the behavioural context in which that information is maintained.

**Significant Statement:** Sound textures, such as rain or fire, are characterised by statistical properties that unfold over time, yet it remains unclear how the brain maintains these representations in working memory. Our findings provide evidence that auditory cortical representations retain information that distinguishes sound textures during working-memory maintenance, while medial temporal lobe representations reflect the behavioural task-related information.

## Introduction

Our listening environments are filled with rich acoustic backgrounds, such as falling rain, crackling fires, and wind. These natural sounds are often described as *auditory textures* because they arise from the overlapping occurrence of many acoustically similar events. A defining characteristic of auditory textures is their approximate stationarity: their statistical properties remain relatively stable over moderate timescales. This stability suggests that texture identity can be represented by summary statistics of acoustic features accumulated over time (McDermott and Simoncelli, 2011; but see Bastug et al., 2026). This is consistent with a broader idea that the auditory system is adapted to efficiently represent the statistical structure of natural sounds (Barlow, 1961; Rieke et al., 1995; Theunissen and Elie, 2014).

McDermott and Simoncelli (2011) demonstrated that a wide range of sound textures can be characterised by summary statistics derived from a model based on the spectral energy and temporal modulation extracted from the peripheral auditory representation. Synthetic sounds generated by matching these statistics to those of natural textures were perceptually similar to the original sounds, suggesting that these statistics capture important aspects of texture identity. Because reliable estimates of such statistics depend on the amount of acoustic information available, McDermott et al. (2013) subsequently examined how stimulus duration influences texture discrimination. They found that discrimination between textures improved as stimulus duration increased over moderate timescales, consistent with the proposal that longer sounds allow more reliable estimation of the statistical properties that distinguish texture categories.

Neurophysiological and neuroimaging studies suggest that statistical properties of sound textures are represented throughout the auditory system, including auditory cortical regions (Zhai et al., 2020; Peng et al., 2024; Mohammadi et al., 2026). However, emerging evidence also implicates the medial temporal lobe (MTL) in the representation and organisation of complex acoustic information (Kumar et al., 2014; Hu et al., 2024; Bianco et al., 2026). The hippocampus, in particular, has been proposed to contribute not only to memory but also to perceptual inference, prediction, and the organisation of information according to contextual demands (Hindy et al., 2016; Stachenfeld et al., 2017). Consistent with a role beyond simple sensory representation, Mohammadi et al. (2026) found increased hippocampal–auditory cortical connectivity as the statistical structure of sound textures became less natural.

Studies of working memory allow the examination of the features of sound that are encoded after initial perception. Previous studies have demonstrated sustained maintenance-period activity in the hippocampus, auditory cortex, and inferior frontal regions when participants maintain auditory information for later comparison (Kumar et al., 2016, 2021). Multivariate activity patterns in the auditory cortex can represent stimulus-specific information during maintenance (Linke et al., 2011; Kumar et al., 2016; Uluç et al., 2018). In the present study, we sought ensemble neural activity and patterns that increase with the duration of stimuli: potentially reflecting the retention of summary statistics or detailed acoustic information. We also investigated the effect of task on retention activity.

Participants performed a texture-discrimination task and a matched loudness-discrimination task using stimuli presented for either 0.1s or 2s. The loudness task allowed us to test effects that were specifically associated with texture processing from more general benefits of increased stimulus duration. Using fMRI, we examined both univariate and multivariate activity patterns during the maintenance period. In the auditory cortex, longer stimuli did not affect activity but enhanced the representation of texture category. In contrast, MTL activity increased for longer stimuli but did not affect the representation of texture category. MTL patterns contained information about the task being performed. Together, these findings reveal distinct representational profiles across auditory cortical and MTL regions during auditory working memory.

## Methods

### Participants

Twenty-five individuals (aged 18-34 years, median 22 years, 8 males, 17 females) with self-reported normal hearing took part. Informed consent was obtained from all participants, who were compensated for their time. The study was approved by the University College London Research Ethics Committee (22805/001).

### Stimuli

Four natural sound texture categories (rain, wind, fire, and insects) were used. Sounds were synthesised using a texture synthesis algorithm (McDermott and Simoncelli, 2011) to produce 5-s signals at an initial sampling rate of 20 kHz. From this synthesised signal, for each trial, a 0.1-or 2-s excerpt was taken from a random starting point within the first 2 s of the signal. The extracted segment was then resampled to 44.1 kHz. Sounds were delivered using a bespoke high-fidelity delivery system based on a manifold of 8 parallel piezoelectric transducers near each ear, connected to a short pneumatic delivery system that achieves high-fidelity presentation of broadband stimuli like the textures here. For all stimuli, an equalisation filter was applied to compensate for the sound delivery system attenuation. Then, root-mean-square was adjusted to a fixed target value of 0.1. A 10 ms cosine ramp (half-cosine onset and offset) was then applied. Subsequently, loudness normalisation was performed using an iterative procedure based on the Glasberg and Moore loudness model (Glasberg and Moore, 2002).

### Paradigm

We presented sounds in a 2× 2 design with Task (texture-type discrimination, *texture* condition, and loudness discrimination, *loudness* condition) and Duration (excerpt with 0.1s and 2s durations) as factors. The experiment consisted of four runs, each containing 64 trials (256 trials in total). There were 32 trials for each task per run, and trials were balanced such that each of the four texture types appeared equally within each task (8 trials per texture).

In each trial, participants performed a delayed match-to-sample task. First, a task instruction cue was displayed for 1.5 s, indicating the task type and whether the participant should attend to “texture” or “loudness” (Fig. 1A). This was followed by a fixation cross while the reference sound was played. Then, after a variable delay (2-6 s), during which the fixation cross remained on screen, participants heard a test sound. Delay intervals were predefined for each run as a vector of 16 values ranging between 2 and 6 s, with more values at shorter and longer delays (mean = 4, std = 1.71). The response prompt was task-dependent: participants were asked either whether the sounds originated from the same texture-type or whether they had the same loudness. Participants had up to 3 s to respond by pressing 1 for same and 2 for different. Trials without a response were recorded as missing. This was followed by an inter-trial interval (1–3 s).

**Figure 1.**
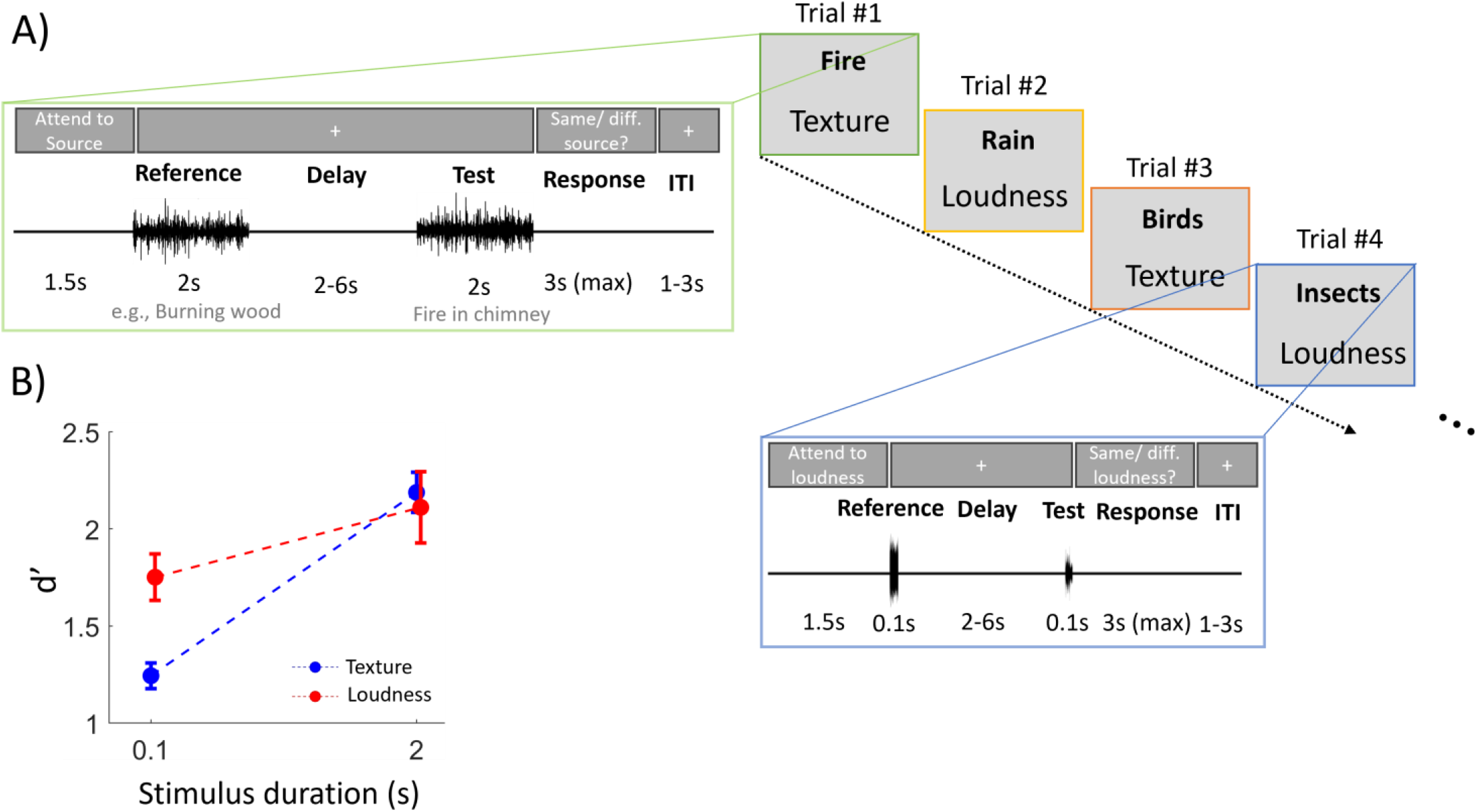
Texture working memory paradigm and discrimination performance. A) Each trial consisted of two sounds, a “reference” and a “test” that either shared the same or differed in texture and loudness. Sounds were presented with either short (0.1s) or long (2s) duration. These sounds were separated by a variable delay period. Four categories of natural texture sounds were used: Birds, Rain, Fire, and Insects. The task instructions were shown on screen with a grey background. B) Discrimination performance (measured as d’). Error bars represent the standard error of the mean. ITI: inter-trial interval.

In the texture task, the texture type for reference and test was either the same or different. When texture types were different, two alternative types were selected for the test sound. For instance, as shown in Fig. 1A, trial 1 has two different sound types (*Burning wood* and *Fire in chimney)* from the same category, fire. The number of trials with the same or different texture types was balanced in each run.

Loudness of stimuli depended on the task condition. In loudness discrimination trials, it was either the same or a different level. When it was the same, both stimuli were normalised either to the standard target (24 sones) or to a lower control level (14 sones), with equal probability. In different loudness trials, one stimulus (reference or test) was normalised to 24 sones while the other to 14 sones. In texture discrimination trials, both reference and test stimuli were normalised to the same target loudness (24 sones).

Psychophysics Toolbox (Brainard, 1997) was used to run the experiment. Before the main fMRI session, participants performed the task outside the scanner room. This consisted of two parts: practice with feedback followed by practice without feedback. In the practice with feedback, 32 trials were presented (one run), and immediate feedback was provided after each response, with a corresponding message (“Correct” or “Incorrect”) displayed for 2 s. This feedback loop was omitted in the practice without feedback. The texture sounds used here had a distinct category from those used in the main fMRI experiment. No feedback was given in the fMRI experiment.

### MRI data acquisition

Data were acquired using a 3-Tesla Siemens MAGNETOM Prisma MR scanner (Siemens Healthcare) with a 32-channel receiver head coil at the Department of Imaging Neuroscience (London, UK). Whole-brain T2*-weighted functional images were acquired using 3D echo-planar imaging (EPI) with a field of view of 192 x 192 x 144 mm, voxel size 3 mm isotropic, 48 partitions, 3D acceleration factor 2, repetition time (TR) 44ms, volume acquisition time 1.056 s, anterior-to-posterior EPI phase encoding, and bandwidth 2298 Hz/Px. Field maps (short echo time (TE) = 10.00 ms, long TE = 12.46 ms) and a whole-brain T1-weighted anatomical image (MPRAGE, field of view 256 x 256 x 176 mm, voxel size 1 mm isotropic, GRAPPA acceleration factor 2, TR = 2530 ms, TE = 3.34 ms) were acquired during the same scanning session.

During scanning sessions, peripheral measurements of subject pulse rate and respiratory signals were made together with scanner slice synchronisation pulses using the Spike2 data acquisition system (Cambridge Electronic Design Limited, Cambridge, UK). The cardiac pulse signal was measured using an MRI-compatible pulse oximeter (Model 8600 F0, Nonin Medical, Inc., Plymouth, MN) attached to the subject’s left index finger. The respiratory signal movement was monitored using a pneumatic belt positioned around the abdomen close to the diaphragm.

A physiological noise model was constructed to account for artefacts related to cardiac and respiratory phase and changes in respiratory volume using an in-house developed MATLAB toolbox (Hutton et al., 2011). This resulted in a total of 14 regressors, which were sampled at a reference slice in each image volume to give a set of values for each time point. The resulting regressors were included as confounds in the first-level analysis for each subject.

To ensure that sounds were presented at a comfortable listening level, we conducted a calibration test inside the MRI scanner with an EPI sequence running, before the main experiment. Participants listened to a continuous sound composed of a mixture of synthesised textures, presented initially at a baseline sound pressure level (SPL) of 80 dB. They were instructed to press a button to gradually increase the volume until they reached a level that they found comfortable, while ensuring that all texture sounds remained clearly audible. The average SPL across participants was 85 dB.

### MRI preprocessing

MRI data were processed in SPM12 (Department of Imaging Neuroscience, UCL). Each participant’s functional images were unwarped using their field map and realigned to the first volume. Functional and anatomical images were coregistered to the mean functional image. For the univariate analyses, the coregistered images were normalised to the SPM12 template (avg305T1) in MNI space, and the functional images were spatially smoothed with a Gaussian kernel with 6 mm full-width at half-maximum (FWHM).

After the data were preprocessed using SPM, the artefact detection (ART) toolbox, implemented in the CONN toolbox (Whitfield-Gabrieli and Nieto-Castanon, 2012), was used to detect outliers from smoothed images. Time points in the series were marked as outliers if the global signal exceeded 3 standard deviations and the movement exceeded 1 mm. Additionally, five covariates generated using the aCmpCorr method (Behzadi et al., 2007), which uses principal component analysis on the measurements made in the white matter (WM) and cerebrospinal fluid (CSF) of each subject’s segmented WM and CSF masks, were included as confounds in the first-level analysis for each subject

### Behavioural data analysis

We measured d’ for each stimulus duration and task type. A repeated-measures ANOVA was used to test the main and interaction effects for Task and Duration factors. Post hoc t-tests were applied to d’ values to test for condition differences, and the tests were corrected for multiple comparisons using the Bonferroni-Holm method.

### Univariate fMRI analysis

First-level general linear model (GLM) analyses were performed using SPM for the whole brain. For each run, a design matrix was constructed that modelled (i) reference and test stimulus presentations for texture and loudness conditions, (ii) maintenance periods split by task and duration, and (iii) ITIs. All regressors were defined using their respective onset times and durations and convolved with the canonical hemodynamic response function. Multiple nuisance regressors were included, consisting of motion parameters, outlier (scrubbing) regressors, physiological recordings and region-of-interest (ROI)-derived WM and CFS signals. A high-pass filter (128 s cutoff) and a first-order autoregressive model were applied to account for low-frequency drift and temporal autocorrelation, respectively. The t-contrasts were defined to test the main effect of duration, the main effect of task and the duration×task interaction effect.

### Anatomical ROIs

Individual T1-weighted images were processed using FreeSurfer’s automated recon-all pipeline to perform anatomical segmentation. The resulting subject-specific cortical parcellations were used to identify auditory cortical regions of interest, including Heschl’s gyrus and planum temporale.

Based on previous literature (Kumar et al., 2016), we defined the medial temporal lobe as our main anatomical ROI comprising the (anterior and posterior) hippocampus, parahippocampal cortex, and entorhinal cortex. Anatomical segmentations were done using the ASHS-PMC-T1 atlas (Xie et al., 2019). The segmentation was done using ITK-SNAP Distributed Segmentation Services.

### Multivariate fMRI analysis

First-level GLMs used the same design as the univariate analysis but were fitted to non-normalised, unsmoothed functional data. Maintenance-period beta images were entered into ROI-based multivariate pattern analyses using The Decoding Toolbox (TDT; Hebart et al., 2015) in MATLAB. Two decoding analyses were performed. Texture-category decoding classified the four sound categories (rain, wind, fire, and insects) separately for each task and stimulus duration (0.1s and 2s) within Heschl’s gyrus (HG) and planum temporale (PT); chance accuracy was 25%. Task decoding classified texture versus loudness trials separately for each duration within the anterior hippocampus, posterior hippocampus, parahippocampal cortex, and entorhinal cortex; chance accuracy was 50%.

For both analyses, a support vector machine implemented in LIBSVM (Chang and Lin, 2011) was used with leave-one-session-out cross-validation, training on three sessions and testing on the remaining session. Feature vectors were z-score normalised before classification. Decoding accuracies were tested against chance using one-tailed *t*-tests. *P*-values were false discovery rate (FDR) corrected for multiple comparisons.

### Code and data availability

Study data are available *here*.

## Results

### Behavioural performance

Behavioural data are shown in Fig. 1B. A repeated-measures ANOVA on *d′* revealed a significant main effect of stimulus duration (*F*(1, 24) = 34.67, *p* < .001), and a significant task × duration interaction (*F*(1, 24) = 18.84, *p* = .0002). The main effect of task was not significant (*F*(1, 24) = 3.07, *p* = .093).

Follow-up pairwise comparisons showed that *d′* was higher for 2s than 0.1s sound for both the texture task (*t*(24) = 8.24, *p* < .001) and the loudness task (*t*(24) = 2.52, *p* = .019). At the 0.1s duration, performance was higher on the loudness task than on the texture task (*t*(24) = 3.65, *p* = .001). In contrast, performance did not differ significantly between tasks at the 2s duration (*t*(24) = 0.55, *p* = .59). Thus, although performance on both tasks benefited from increased stimulus duration, the improvement was substantially greater for texture discrimination.

### Neural activity during maintenance

Whole-brain voxel-wise GLM analyses were performed for the maintenance period. There was no significant main effect of task and no significant task × duration interaction. We observed a significant main effect of stimulus duration in the medial temporal lobe (MTL) at an uncorrected threshold of *p* < .001 (Fig. 2, top). This effect survived after FWER correction within the MTL volume of interest (*p* < .05; Fig. 2, bottom), with a peak at MNI coordinates *x* = 24, *y* = −12, *z* = −16, *t*(24) = 6.96. This result shows MTL activity during the maintenance period varied as a function of the duration of the preceding auditory stimulus, irrespective of task.

**Figure 2.**
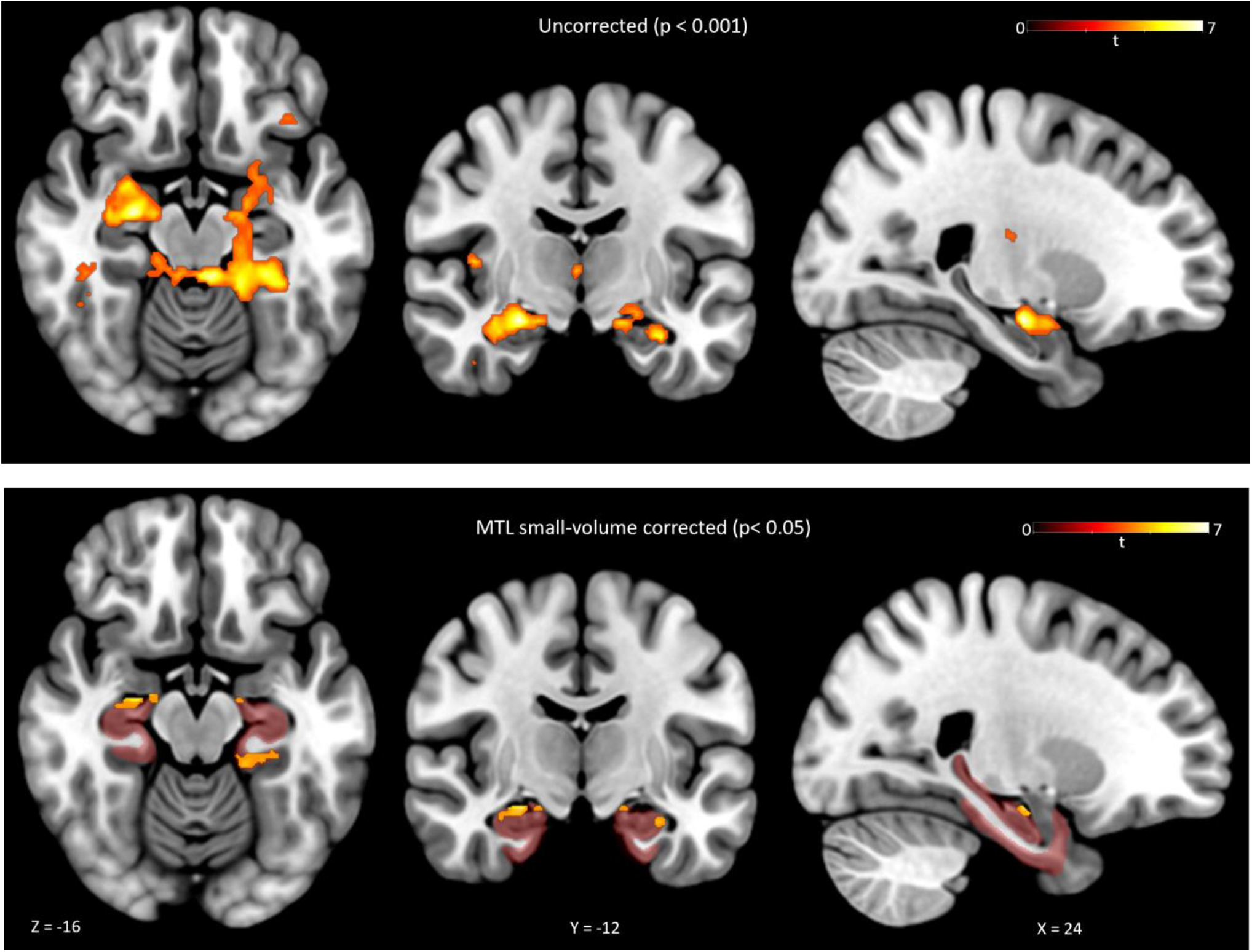
Univariate analysis of BOLD signal. Top panel: the effect of stimulus duration (long > short) on maintenance-period BOLD responses with an uncorrected (*p* < 0.001) threshold. Bottom panel, the duration effect for the medial temporal lobe (MTL) region (*p* < 0.05, small-volume corrected). ROI is shown with red shading.

### Decoding of sound texture category from auditory cortex

A whole-brain searchlight analysis revealed regions in the auditory cortex in which texture-category decoding during maintenance was higher for 2s than 0.1s stimuli at the uncorrected threshold of *p* < 0.001 (Fig. 3, left panel). To characterise this effect further, ROI-based decoding analyses were performed separately in HG and PT for each duration and task.

**Figure 3.**
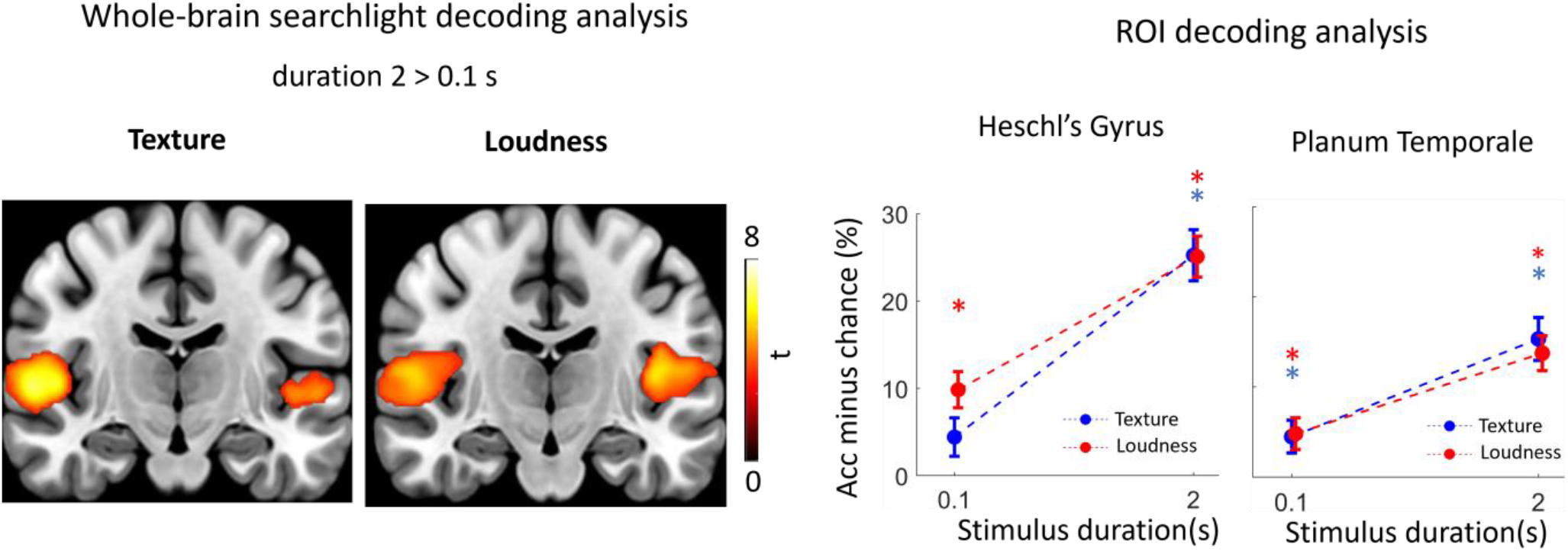
Sound texture category decoding during the maintenance period. Left, whole-brain searchlight analysis showing regions with significant decoding differences between stimulus durations (*p* < 0.001, uncorrected). Right, ROI-based decoding in Heschl’s gyrus and planum temporale. Asterisks indicate decoding accuracy significantly above chance (25%; *p* < 0.001, FDR-corrected), with bars coloured according to task (texture or loudness).

Texture-category decoding was significantly above chance (25%) in all conditions (*p*_FDR_ < 0.001) except the 0.1s texture task in HG (*p*_FDR_ > 0.05). In both HG and PT, decoding performance showed a significant main effect of stimulus duration (*p* < 0.001), with higher decoding accuracy for 2s than 0.1s stimuli. There was no significant main effect of task and no significant task × duration interaction. These results show texture-category decoding in auditory cortical activity patterns during the maintenance period and that the decoding performance increased with stimulus duration.

### Decoding of task context from MTL regions

As shown above in the searchlight decoding analysis, no evidence for texture-category decoding was observed in the MTL. We next tested whether the task being performed could be decoded from activity patterns in MTL regions during the maintenance period. For the 2-s condition, task decoding was significantly above chance in the posterior hippocampus, *t*(24) = 2.80, *p*_FDR_ = .029, and in the parahippocampal cortex (PHC), *t*(24) = 4.08, *p*_FDR_ = .002 (Fig. 4). We then tested whether accuracy was higher for longer compared with shorter stimulus durations in each region and found a significant duration effect in the PHC, *t*(24) = 2.46, *p*_FDR_ = .04. These findings indicate that, following longer stimuli, distributed activity patterns in MTL regions contained information distinguishing whether participants were maintaining auditory information for texture or loudness discrimination, with a significant increase in task-decoding accuracy for longer stimuli in the PHC.

**Figure 4.**
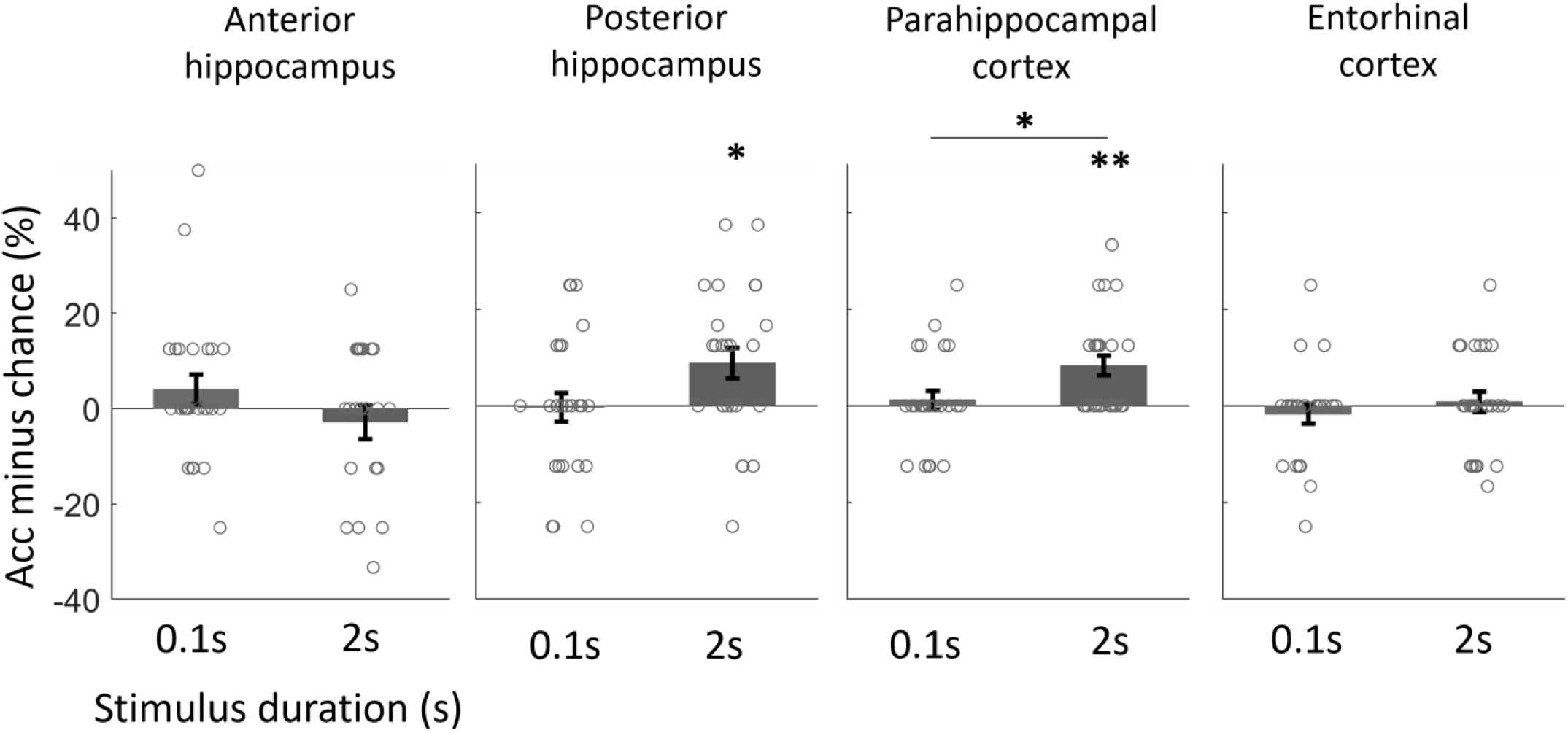
Percentage accuracy of pattern classification (texture vs loudness) during the maintenance period using multivoxel pattern analyses. The mean accuracy minus chance (50%) classifier performance is shown (SEM across subjects, individual subject data shown with open circles). Accuracy (Acc) was above chance level in the posterior hippocampus (* *p* = 0.029, FDR-corrected) and parahippocampal cortex (** *p*_*FDR*_ = 0.002) during the long stimulus duration. Greater accuracy for longer versus shorter duration is observed in the parahippocampal cortex (*p*_*FDR*_ = 0.04).

## Discussion

The present findings show that the duration effect of an auditory sound texture on texture perception occurs at both behavioural and neural levels. Behaviourally, increasing stimulus duration from 0.1 s to 2 s improved performance in both the texture and loudness tasks, but the benefit was substantially greater for texture discrimination. This pattern is consistent with previous work suggesting that auditory texture discrimination benefits from accumulating summary statistics over time (McDermott and Simoncelli, 2011; McDermott et al., 2013). However, increasing duration could also provide additional opportunities to sample and encode local acoustic features that distinguish textures (Bastug et al., 2026). The improvement in loudness discrimination may partly reflect the well-established phenomenon of temporal loudness summation (Zwislocki, 1969). Nevertheless, the larger improvement for texture discrimination suggests that texture judgments may benefit more strongly from the accumulation of acoustic information over time.

Animal studies have demonstrated that neurons in the inferior colliculus are sensitive to both low-and higher-order statistical properties of natural sound textures, with this sensitivity reflected in firing-rate responses and correlation of neural population activity (Sadeghi et al., 2019; Zhai et al., 2020; Mishra et al., 2021; Peng et al., 2024). A recent human study has shown that auditory cortical regions are also sensitive to the statistics of natural sound textures (Mohammadi et al., 2026). In the present study, we decoded texture category from multivariate activity patterns in Heschl’s gyrus and Planum Temporale during the maintenance period, and decoding performance was higher for 2s than for 0.1 s sounds. Also, we did not observe increased ensemble response for longer sounds than for shorter ones during maintenance. Longer sound durations may therefore yield more discriminable neural representations of texture category despite the same level of activity. This is consistent with previous evidence that natural sound category and spectrotemporal properties are represented in distributed patterns across the auditory cortex (Staeren et al., 2009; Santoro et al., 2014). We also observed significant texture decoding in the loudness task. Note that in this task the participants heard sound textures but had to maintain loudness information. This decoding result therefore suggests that auditory cortical representations of texture category were not restricted to conditions in which texture identity was explicitly relevant for behaviour. Longer duration may therefore enhance the discriminability of texture-related information in the auditory cortex even when texture identity is not explicitly task relevant.

We observed that MTL activity during maintenance increased with the duration of the sound irrespective of task. The duration effect therefore cannot readily be attributed to selective engagement of the MTL during texture discrimination. A potential concern might be that the effect reflects residual haemodynamic activity evoked by the longer stimulus, given the temporally extended nature of the BOLD response (Boynton et al., 1996; Robson et al., 1998). However, the absence of a corresponding duration effect in auditory cortex argues against a simple global consequence of longer stimulus presentation. Nevertheless, the MTL duration effect is compatible with greater engagement of mnemonic processes following longer auditory input, consistent with previous evidence for sustained hippocampal involvement during auditory working-memory maintenance (Kumar et al., 2016).

Multivariate analyses for MTL regions during maintenance showed that activity patterns in the posterior hippocampus and parahippocampal cortex distinguished the texture and loudness task, whereas we found no evidence that the same regions discriminated texture category. This may indicate a greater sensitivity of MTL representations to behavioural context than to texture identity. This may also reflect differences in the cognitive demands of the two tasks, with texture discrimination potentially requiring the maintenance of more detailed acoustic information, whereas loudness discrimination may rely on a simpler representation of overall intensity. These findings are consistent with evidence that hippocampal representations are shaped by attentional state, task structure, and current behavioural goals (Aly and Turk-Browne, 2016; Córdova et al., 2019; Berens and Bird, 2022; Crivelli-Decker et al., 2023). This contrasts with the auditory cortex, where texture-category information was decodable across tasks and became more discriminable following longer stimulus presentations.

## Conclusion

Together, our findings show a sound-duration effect on MTL activity and on texture-category representation in the auditory cortex during maintenance, regardless of task instruction. However, task representation can be decoded in the posterior hippocampus and parahippocampal cortex. The results suggest that longer texture duration supports more discriminable representations of sound category and highlight a role for the MTL in representing task-related context.

## Acknowledgements

This work was supported by the Medical Research Council, United Kingdom (grant number MR/T032553/1). We thank the Imaging Support team at the UCL Department of Imaging Neuroscience.

## References

Aly M, Turk-Browne NB (2016) Attention Stabilizes Representations in the Human Hippocampus. Cereb Cortex 26:783–796 Available at: https://pubmed.ncbi.nlm.nih.gov/25766839/.

Barlow HB (1961) Possible Principles Underlying the Transformations of Sensory Messages. Sensory Communication:217–233.

Bastug B, Rajendran VG, Bianco R, Agus T, Chait M, Pressnitzer D (2026) Memory for repeated auditory textures. Cognition 268.

Behzadi Y, Restom K, Liau J, Liu TT (2007) A component based noise correction method (CompCor) for BOLD and perfusion based fMRI. Neuroimage 37:90–101 Available at: https://pubmed.ncbi.nlm.nih.gov/17560126/.

Berens SC, Bird CM (2022) Hippocampal and medial prefrontal cortices encode structural task representations following progressive and interleaved training schedules. PLoS Comput Biol 18:e1010566 Available at: https://journals.plos.org/ploscompbiol/article?id=10.1371/journal.pcbi.1010566 [Accessed August 12, 2026].

Bianco R, Magami K, Pearce M, Chait M (2026) Discovery, Interruption, and Updating of Auditory Regularities in Memory: Evidence from Low-Frequency Brain Dynamics in Human MEG. J Neurosci 46 Available at: https://pubmed.ncbi.nlm.nih.gov/41266245/.

Boynton GM, Engel SA, Glover GH, Heeger DJ (1996) Linear systems analysis of functional magnetic resonance imaging in human V1. J Neurosci 16:4207–4221 Available at: https://pubmed.ncbi.nlm.nih.gov/8753882/ [Accessed August 12, 2026].

Brainard DH (1997) The psychophysics toolbox. Spat Vis:433–436.

Chang CC, Lin CJ (2011) LIBSVM: A Library for support vector machines. ACM Trans Intell Syst Technol 2:27 Available at: https://dl.acm.org/doi/pdf/10.1145/1961189.1961199.

Córdova NI, Turk-Browne NB, Aly M (2019) Focusing on what matters: Modulation of the human hippocampus by relational attention. Hippocampus 29:1025–1037 Available at:/doi/pdf/10.1002/hipo.23082 [Accessed August 12, 2026].

Crivelli-Decker J, Clarke A, Park SA, Huffman DJ, Boorman ED, Ranganath C (2023) Goal-oriented representations in the human hippocampus during planning and navigation. Nature Communications 2023 14:1 14:2946–Available at: https://www.nature.com/articles/s41467-023-35967-6 [Accessed August 12, 2026].

Glasberg BR, Moore BCJ (2002) A Model of Loudness Applicable to Time-Varying Sounds. Available at: https://api.semanticscholar.org/CorpusID:52265560.

Hebart MN, Görgen K, Haynes JD (2015) The decoding toolbox (TDT): A versatile software package for multivariate analyses of functional imaging data. Front Neuroinform 8:108528 Available at: www.frontiersin.org.

Hindy NC, Ng FY, Turk-Browne NB (2016) Linking pattern completion in the hippocampus to predictive coding in visual cortex. Nat Neurosci 19:665–667 Available at: https://pubmed.ncbi.nlm.nih.gov/27065363/.

Hu M, Bianco R, Hidalgo AR, Chait M (2024) Concurrent Encoding of Sequence Predictability and Event-Evoked Prediction Error in Unfolding Auditory Patterns. Journal of Neuroscience 44 Available at: https://www.jneurosci.org/content/44/14/e1894232024.

Hutton C, Josephs O, Stadler J, Featherstone E, Reid A, Speck O, Bernarding J, Weiskopf N (2011) The impact of physiological noise correction on fMRI at 7 T. Neuroimage 57:101–112 Available at: https://www.sciencedirect.com/science/article/pii/S105381191100396X?via%3Dihub.

Kumar S, Bonnici HM, Teki S, Agus TR, Pressnitzer D, Maguire EA, Griffiths TD (2014) Representations of specific acoustic patterns in the auditory cortex and hippocampus. Proceedings of the Royal Society B: Biological Sciences 281:20141000 Available at: https://royalsocietypublishing.org/rspb/article/281/1791/20141000/77261/Representations-of-specific-acoustic-patterns-in.

Kumar S, Gander PE, Berger JI, Billig AJ, Nourski K V, Oya H, Kawasaki H, Howard MA, Griffiths TD (2021) Corrigendum to “Oscillatory correlates of auditory working memory examined with human electrocorticography” [Neuropsychologia 150 (2021) 107691]. Neuropsychologia 159:107950 Available at: http://www.ncbi.nlm.nih.gov/pubmed/34246979.

Kumar S, Joseph S, Gander PE, Barascud N, Halpern AR, Griffiths TD (2016) A Brain System for Auditory Working Memory. Journal of Neuroscience 36:4492–4505 Available at: https://www.jneurosci.org/content/36/16/4492.

Linke AC, Vicente-Grabovetsky A, Cusack R (2011) Stimulus-specific suppression preserves information in auditory short-term memory. Proc Natl Acad Sci U S A 108:12961–12966 Available at: https://pubmed.ncbi.nlm.nih.gov/21768383/ [Accessed September 29, 2026].

McDermott JH, Simoncelli EP (2011) Sound texture perception via statistics of the auditory periphery: Evidence from sound synthesis. Neuron 71:926–940.

Mishra AP, Peng F, Li K, Harper NS, Schnupp JWH (2021) Sensitivity of neural responses in the inferior colliculus to statistical features of sound textures. Hear Res 412 Available at: https://pubmed.ncbi.nlm.nih.gov/34739889/ [Accessed October 27, 2026].

Mohammadi Y, Billig AJ, Berger JI, Griffiths TD (2026) Brain Representations of Natural Sound Statistics. Journal of Neuroscience:e1204252026 Available at: https://www.jneurosci.org/content/early/2026/05/11/JNEUROSCI.1204-25.2026 [Accessed May 19, 2026].

Peng F, Harper NS, Mishra AP, Auksztulewicz R, Schnupp JWH (2024) Dissociable Roles of the Auditory Midbrain and Cortex in Processing the Statistical Features of Natural Sound Textures. Journal of Neuroscience 44 Available at: https://www.jneurosci.org/content/44/10/e1115232023 [Accessed April 19, 2025].

Rieke F, Bodnar DA, Bialek W (1995) Naturalistic stimuli increase the rate and efficiency of information transmission by primary auditory afferents. Proceedings of the Royal Society B: Biological Sciences 262:259–265 Available at: 10.1098/rspb.1995.0204.

Robson MD, Dorosz JL, Gore JC (1998) Measurements of the temporal fMRI response of the human auditory cortex to trains of tones. Neuroimage 7:185–198 Available at: https://pubmed.ncbi.nlm.nih.gov/9597660/.

Sadeghi M, Zhai X, Stevenson IH, Escabí MA (2019) A neural ensemble correlation code for sound category identification. PLoS Biol 17:e3000449 Available at: https://journals.plos.org/plosbiology/article?id=10.1371/journal.pbio.3000449 [Accessed October 27, 2026].

Santoro R, Moerel M, De Martino F, Goebel R, Ugurbil K, Yacoub E, Formisano E (2014) Encoding of Natural Sounds at Multiple Spectral and Temporal Resolutions in the Human Auditory Cortex. PLoS Comput Biol 10:e1003412 Available at: https://journals.plos.org/ploscompbiol/article?id=10.1371/journal.pcbi.1003412 [Accessed August 12, 2026].

Stachenfeld KL, Botvinick MM, Gershman SJ (2017) The hippocampus as a predictive map. Nat Neurosci 20:1643–1653 Available at: https://pubmed.ncbi.nlm.nih.gov/28967910/.

Staeren N, Renvall H, De Martino F, Goebel R, Formisano E (2009) Sound Categories Are Represented as Distributed Patterns in the Human Auditory Cortex. Current Biology 19:498–502 Available at: https://www.sciencedirect.com/science/article/pii/S0960982209007404?via%3Dihub.

Theunissen FE, Elie JE (2014) Neural processing of natural sounds. Nature Reviews Neuroscience 2014 15:6 15:355–366 Available at: https://www.nature.com/articles/nrn3731.

Uluç I, Schmidt TT, Wu Y hao, Blankenburg F (2018) Content-specific codes of parametric auditory working memory in humans. Neuroimage 183:254–262 Available at: https://pubmed.ncbi.nlm.nih.gov/30107259/ [Accessed September 29, 2026].

Whitfield-Gabrieli S, Nieto-Castanon A (2012) Conn: A Functional Connectivity Toolbox for Correlated and Anticorrelated Brain Networks. Brain Connect 2:125–141 Available at:/doi/pdf/10.1089/brain.2012.0073?download=true.

Xie L, Wisse LEM, Pluta J, de Flores R, Piskin V, Manjón J V., Wang H, Das SR, Ding SL, Wolk DA, Yushkevich PA (2019) Automated segmentation of medial temporal lobe subregions on in vivo T1-weighted MRI in early stages of Alzheimer’s disease. Hum Brain Mapp 40:3431 Available at: https://pmc.ncbi.nlm.nih.gov/articles/PMC6697377/.

Zhai X, Khatami F, Sadeghi M, He F, Read HL, Stevenson IH, Escabí MA (2020) Distinct neural ensemble response statistics are associated with recognition and discrimination of natural sound textures. Proc Natl Acad Sci U S A 117:31482–31493 Available at: https://pubmed.ncbi.nlm.nih.gov/33219122/ [Accessed April 19, 2025].

Zwislocki JJ (1969) Temporal Summation of Loudness: An Analysis. J Acoust Soc Am 46:431–441 Available at:/asa/jasa/article/46/2B/431/644404/Temporal-Summation-of-Loudness-An-Analysis [Accessed November 5, 2026].

